# Large prospective losses lead to sub-optimal sensorimotor decisions in humans

**DOI:** 10.1101/406439

**Authors:** Tyler J. Adkins, Richard L. Lewis, Taraz G. Lee

## Abstract

The rationality of human behavior has been a major problem in philosophy for centuries. The pioneering work of Kahneman and Tversky provides strong evidence that people are not rational. Recent work in psychophysics argues that incentivized sensorimotor decisions (such as deciding where to reach to get a reward) maximizes expected gain, suggesting that it may be impervious to cognitive biases and heuristics. We rigorously tested this hypothesis using multiple experiments and multiple computational models. We obtained strong evidence that people deviated from the objectively rational strategy when potential losses were large. They instead appeared to follow a strategy in which they simplify the decision problem and satisfice rather than optimize. This work is consistent with the framework known as bounded rationality, according to which people behave rationally given their computational limitations.

## Introduction

The rationality of human behavior has been a key problem in philosophy and psychology for centuries and remains so today. Broadly, rationality refers to the conformity of thought and action to reason. In thought, this amounts to forming justified beliefs; in action, this amounts to adopting suitable means to ends. Setting aside issues about what makes something “justified” or “suitable”, it is prima facie intuitive that humans are rational. However, research in behavioral economics has provided evidence that human behavior deviates from what would be expected of perfectly rational utility-maximizing agents(Tversky and Kahneman, 1985, 1979). A prominent example is the work of Kahneman and Tversky which catalogues a rich array of cognitive biases and heuristics that influence human choice behavior, often in detrimental ways(Tversky and Khaneman, 1974). This research has had a tremendous influence on discourse about human rationality and has provided substantial support for the view that humans are fundamentally not rational.

Notwithstanding the doctrine of behavioral economics, many prominent computational models across the subfields of psychology suggest that the human brain continually performs inferences and actions that are optimal in that they minimize costs or maximize gains. Optimization models have proved useful for explaining human perception (Bastos et al., 2012; Feldman and Friston, 2010; Friston, 2010; Knill and Richards, 1996; Ma et al., 2006), cognition (Baker et al., 2011; Dayan and Daw, 2008; Griffiths et al., 2008; Howes et al., 2016; Lewis et al., 2014; Pezzulo et al., 2018; Tauber et al., 2017), and action (Braun et al., 2011; Brown et al., 2011; Diedrichsen et al., 2010; Friston, 2011, 2010; Körding and Wolpert, 2006, 2004; Manohar et al., 2015; Neyedli and Welsh, 2013); however, for a discussion of systematic sub-optimality in human decision-making see (Rahnev and Denison, 2018). One especially compelling demonstration of optimal human behavior is found in studies using a psychophysics task created by Julia Trommershauser and colleagues (Trommershäuser et al., 2008, 2006, 2005, 2003). In these studies, subjects perform rapid reaching movements to small rewarding targets while avoiding overlapping loss regions. This task was designed to be conceptually equivalent to more classical decision-making tasks in behavioral economics in which people display biases such as loss aversion and risk aversion. A key finding from this work is that people shift their aim-points away from the loss regions when losses are larger. The most prominent model of this phenomenon is a utility-maximizing statistical decision model. According to this model, people select aimpoints that maximize expected utility, where expected utility is computed using information about outcome probabilities and outcome values, where outcome probabilities are determined using an internal estimate of outcome variance. This work suggests that utility-maximization may be a fundamental principle of sensorimotor control in humans.

In this paradigm, subjects are presented with perceptual targets and financial incentives, then they plan and execute rapid reaching movements towards these targets. While these adaptations may be optimal in some situations, we suspect that there are situations in which people will deviate systematically from the optimal strategy. In particular, we hypothesize that when payoffs are large, subjects may experience too much motivation, leading them to paradoxically fail. Indeed, choking under pressure in sensorimotor and decision-making tasks is well documented in the literature (Chib et al., 2014; DeCaro et al., 2011; Englert and Oudejans, 2014; Gray, 2004; Kinrade et al., 2010; Lee and Grafton, 2015; Oudejans et al., 2011). According to one prominent account of choking, it occurs when pressure leads to excessive top-down cognitive influences on performance (DeCaro et al., 2011; Englert and Oudejans, 2014; Lee and Grafton, 2015; Oudejans et al., 2011; Snyder and Logan, 2013; Yu, 2015). We expect that this increased reliance on cognition will make task performance susceptible to cognitive biases and heuristics.

We administered an incentivized reaching task to 33 healthy human subjects across two experiments to test whether human sensorimotor control maximizes expected gain, or instead deviates from optimal because of deleterious interactions between motivation and cognition. Consistent with prior work, we found that subjects adapted their reaching movements to payoff information, shifting their aimpoints away from the loss region when losses were large. Formal model comparison revealed that behavior was described best by the optimal model when the losses and gains were of equal magnitude, suggesting that visuomotor decision making may be optimal in medium stakes situations. However, when the losses were large, subjects’ behavior deviated substantially from the optimal model’s predictions, and was described better by a simple heuristic model. Subjects’ visuomotor adaptations to large losses were consistently less-than-optimal, suggesting that subjects are not following the optimal strategy. Overall, our results suggest that human visuomotor control is sensitive to payoffs but not strictly optimal. Instead, when the stakes are high, people appear to abandon the optimal task strategy in favor of a heuristic strategy that is simpler and more stable across people and payoffs.

## Results

### Experiment 1

17 healthy human subjects performed a modified version of the incentivized reaching task used in prior work (Figure 1a). This task was designed to generate heightened motivational pressure, which we hypothesize will make subjects more likely to deviate from the optimal strategy. The task featured varying distances between the origins of gain and loss circles (32px or 44px), and a wide range of payoff conditions ([-$0,+$1], [-$1,+$1], [-$3,+$3], [-$5,+$1], [-$15,+$3]). We classify these payoff conditions based on the loss ratio (0:1, −1:1, −5:1) and the payoff multiplier (1x, 3x). Task conditions varied trial-wise and monetary earnings were determined by the subject’s performance on a trial selected at random at the end of the experiment. Subjects started with a training session that included hundreds of reaches to random dots, as well as a few blocks of practice with each task condition. Subjects returned two days later to perform 10 blocks of 60 trials with opportunities for monetary gain and loss. We measured participants reaction time, movement time, and their reach endpoints. After the completing the test session, subjects performed a separate economic choice task to assess their trait-level loss aversion (Tom et al., 2007).

**Figure 1.**
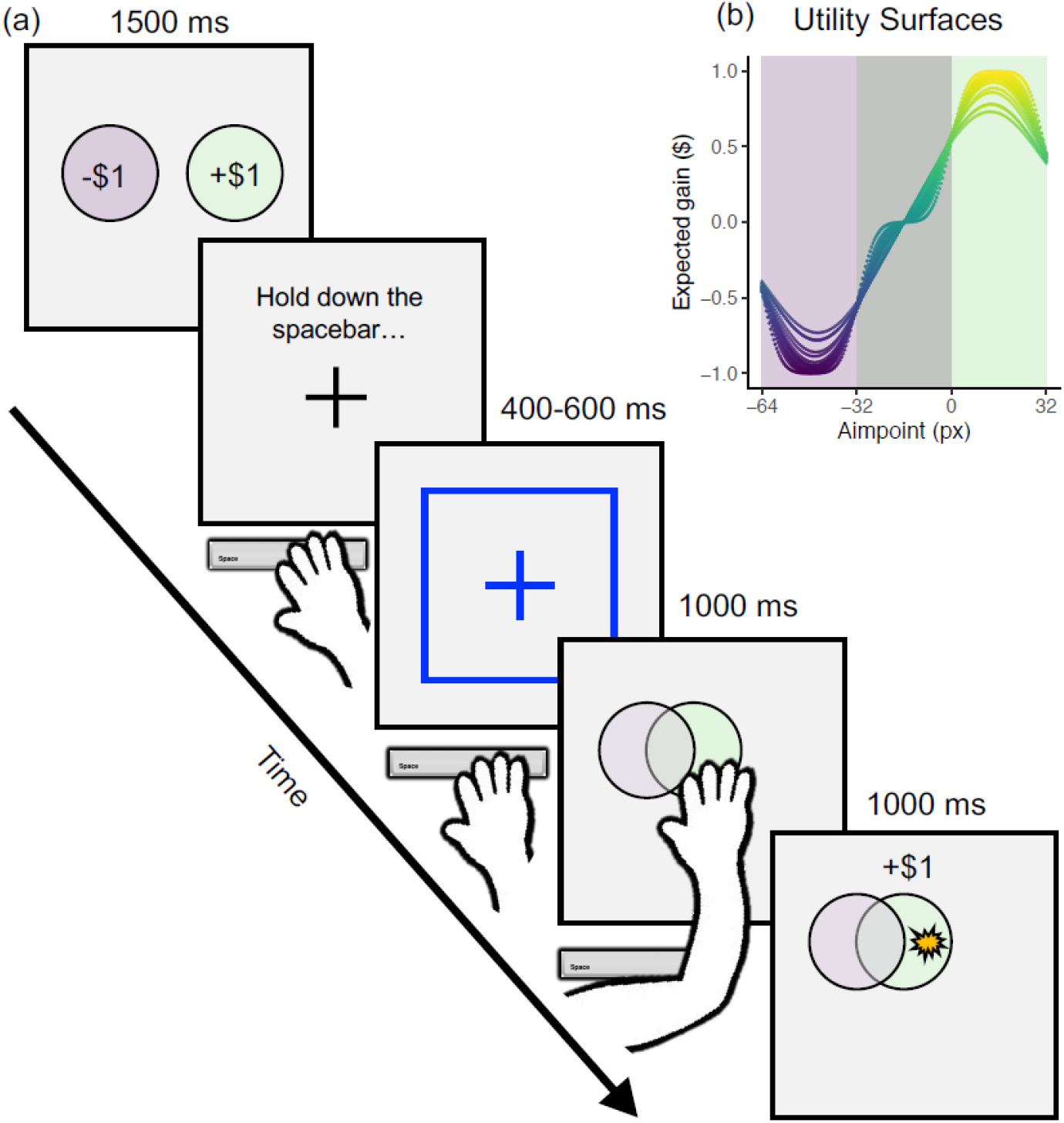
Example trial and utility surface. (a) Participants are first presented with information about the potential payoffs. Next, they initiate the trial by pressing and holding down the spacebar. A fixation cross is presented for a variable delay interval. Next, the stimuli are presented, and the participant releases the spacebar and reaches out to touch the stimulus. They are shown feedback at the end of each trial indicating the location they touched and the payoff they received. (b) Using the optimal model, we can estimate the expected earnings associated with each possible aimpoint. Here we show the shape of this utility surface over a range of x-coordinates and participants (i.e., training variances).

#### Reaching behavior was sensitive to financial stakes and target size

We fit a multilevel gaussian regression model to subjects’ reach endpoints. The model included categorical fixed effects of loss ratio (−1:1 vs −5:1), payoff multiplier (1x vs 3x), and separation distance between loss and gain regions (32px vs 44px), as well as continuous fixed effects of training variance and loss aversion, both of which are constant within subject. If reaching behavior is guided by the optimal strategy, then it will be affected by loss ratio and training variance but not loss aversion or payoff multiplier. We used Bayesian parameter estimation with weakly informative priors (β ~ *N*(0,1)) to estimate the effects of interest. The outcome variable and continuous predictors were mean-centered and scaled to have standard deviations of 0.5 (Gelman, 2008; Gelman et al., 2008). This transformation ensures that our priors assign high probability to small effects (especially, zero) and low probability to large effects. It also ensures that all effects on a common scale, enabling easy comparisons between them. Our model included fixed effects of separation distance, loss ratio, payoff multiplier, loss-aversion, and training variance, as well as random intercepts for each subject. We report the posterior median (β), 89% highest density intervals (HDI) (Kruschke, 2014), and Probability of Direction (pd)(Makowski et al., 2019). Figure 2a shows posterior probability densities over sizes of our effects of interest, Figure 2b shows predicted marginal effects of interest, and Figure 2c shows posterior predictive checks against observed means and standard deviations. We excluded data from the zero-penalty trials because there was no 3x multiplier variant for these trials.

**Figure 2.**
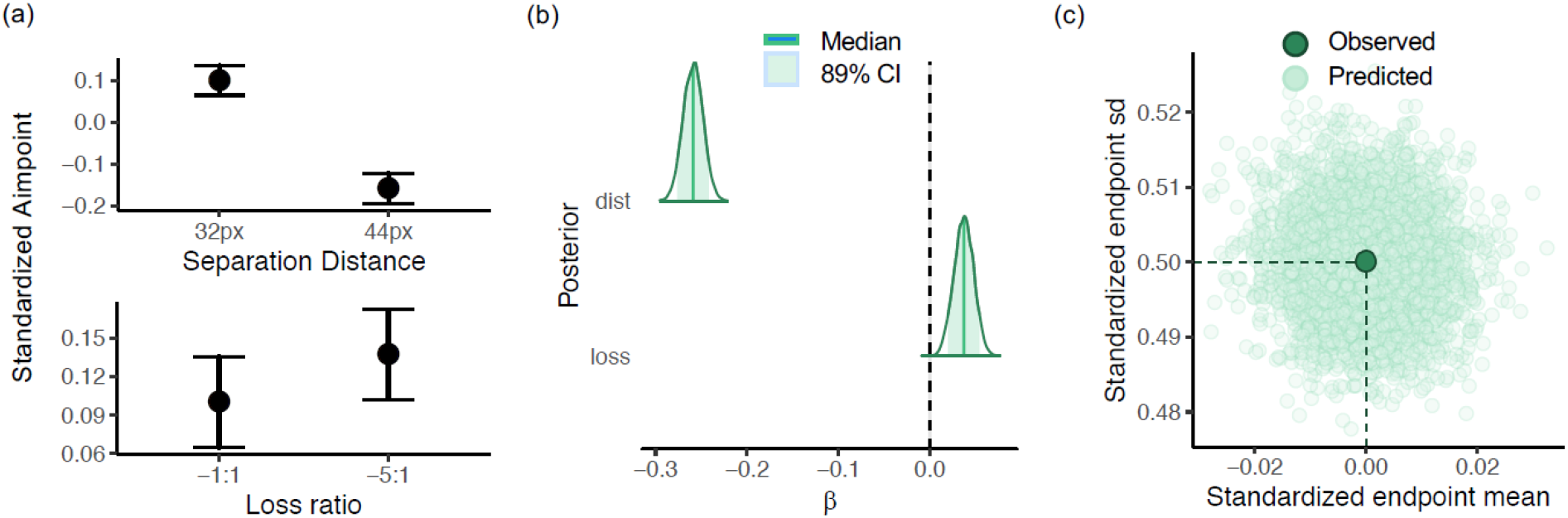
Results from a Bayesian multilevel regression model of horizontal endpoint. (a) Marginal effects of separation distance and loss-to-gain ratio on mean horizontal endpoint. People aimed closer to the loss region when the distance between the circles was greater, and they aimed a bit further from the loss region when the loss ratio was high. (b) Posterior parameter estimates for the effects shown in (a). The shape and location of these posteriors show that sign and magnitude errors are unlikely. (c) Recovery of the empirical grand endpoint mean and sd from 4000 posterior draws.

We found strong evidence for a small (8%sd) positive effect of loss ratio on reach endpoint, suggesting that our subjects reached further from the origin when the ratio of loss to gain was high compared to when it was low (β = 0.04, HDI = [0.02, 0.05], pd = 1.0). We also found strong evidence for a large (50%sd) negative effect of separation distance on reaching endpoint, suggesting that our participants aimed closer to the origin when the separation distance was large compared to when it was small (β = - 0.25, HDI = [-0.28, −0.24], pd = 1.0). We found strong evidence for a very small (2%sd) positive effect of payoff multiplier on reach endpoint, but this effect may be negligibly small (β = 0.02, HDI = [0.00, 0.04], pd = 0.97). We found weak evidence for a small (12%sd) negative effect of loss aversion (β = −0.06, HDI = [-0.12, 0.00], pd = 0.94) and weak evidence for a very small (4%sd) negative effect of training variance (β = −0.02, HDI = [-0.12, 0.00], pd = 0.94), but we refrain from drawing inferences from these estimates given their high posterior uncertainty. In sum, this analysis revealed that human reaching behavior was highly sensitive to visuospatial information and minimally sensitive to financial stakes.

#### Optimal and suboptimal models of reaching behavior

The analysis above showed that reaching behavior in the present task had optimal features, such as sensitivity to loss ratio and training variance, as well as suboptimal features, such as sensitivity to payoff multiplier and loss aversion. We therefore considered multiple computational models of task performance that instantiated different combinations of these features. The first model that we considered has been used in prior work and we refer to it as the optimal model. This model states that a person selects reach aimpoints that maximize expected utility conditioned on an internal estimate of their own reach variance. We implement this model by estimating expected utility surfaces over the stimulus space for each subject and payoff condition and selecting the points that maximize these surfaces (Figure 1b). The shape of the utility surface is determined by the loss-to-gain ratio and the subject’s reach variance estimate, which we estimate from performance in training (Figure 1b). When the loss ratio is high, the peaks of the utility surfaces are shifted away from the loss region suggesting participants are somewhat sensitive to loss ratio (Figure 3a). As for the effect of variance, the utility surfaces for low variance subjects are more extreme (higher peaks, lower valleys), but the maxima of these surfaces are closer to the loss region, compared to the utility surfaces for high variance subjects.

**Figure 3.**
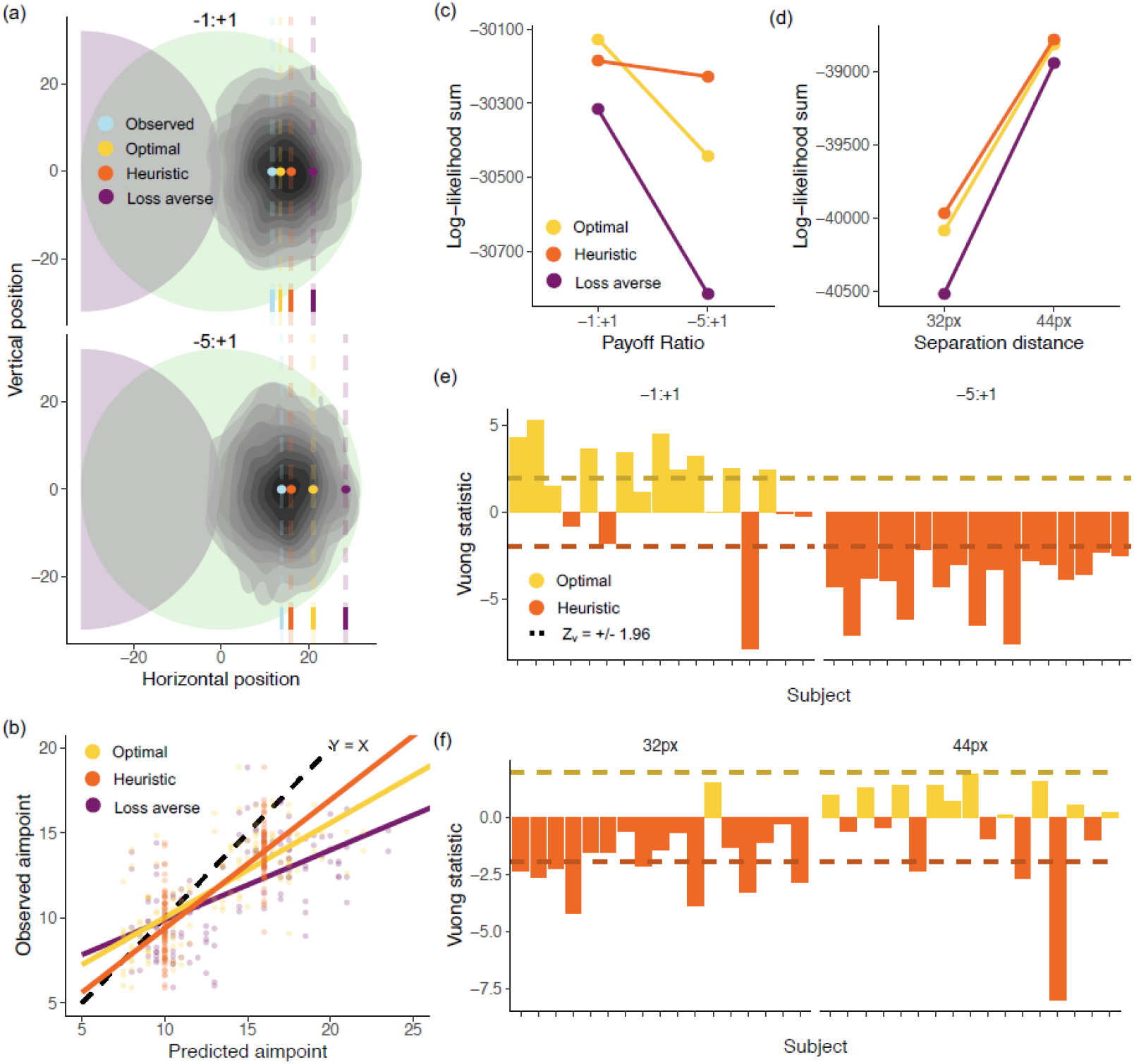
Computational modeling results for Experiment 1. (a) Endpoint distribution and average model predictions for the two loss-ratio conditions. The observed mean endpoint (blue dot) does not move much between the two payoff conditions, similar to the heuristic aimpoint (orange). The optimal aimpoint (yellow) shifts away from the loss region when the loss ratio increases, and this effect is exaggerated for the loss-averse model (purple). This plot is for illustrative purposes, so we show only subjects with above average endpoint variance and above average loss-aversion, to ensure the predictions of the models are visibly distinct. (b) Model-predicted aimpoints vs observed aimpoints. Points on the Y = X line represent perfect correspondence between the model and the data. Most people shift less than predicted by the models, but the heuristic predictions provide the best match to the observed data. There is a separate point for each combination of subject, condition and model. (c) Summed log-likelihoods of the data under each model in each payoff condition—greater log-likelihood indicates better fit. The optimal model fits the best in the −1:1 condition by a small amount, and the heuristic model fits best in the −5:1 condition by a greater amount. It also seems that all three models fit worse in in the higher stakes condition. (d) Log-likelihoods for each spatial condition. It appears that the models fit better in the wide spatial condition. (e) Vuong’s statistic supports likelihood-ratio tests for comparing models. Here we show the results of Vuong’s test comparing the optimal and heuristic models, separately for each subject and payoff condition. These results show that the group-level patterns hold for most individual subjects. (f) Vuong’s statistics for subject-level heuristic vs optimal comparisons for each spatial condition.

We refer to our second model as the loss-averse model. The only difference between this model and the optimal model is that the loss magnitudes were given outsized weight when estimating the utility surfaces. The losses were weighted according to a subject’s trait-level loss aversion that was measured in a separate choice task. This model assumes that people still maximize utility, but the utility is subjective and biased. The overweight losses decrease the utilities of nearby aimpoints and shifts the utility maxima away from the loss region (Figure 3a).

The final model that we considered is called the heuristic model. This model states that people follow a simple intuitive strategy that produces satisfactory albeit strictly suboptimal outcomes. The idea behind this model is that people tend to simplify difficult problems to make them more computationally tractable. In other words, when faced with vast decision-spaces, people tend to satisfice rather than optimize (Simon, 1955).. Therefore, the heuristic model states that people simplify the incentivized reaching task by maximize the probability of hitting the pure-reward region, while ignoring the vertical (less relevant) spatial dimension. In the present task, this strategy leads to aimpoints that depend only on (1) the presence/absence of a potential loss, and (2) the horizontal spatial configuration of the stimuli. When there is a potential loss, the heuristic agent aims at the midpoint of ‘pure gain’ portion of the midline axis (Figure 3a), otherwise the heuristic agent aims at the center of the gain circle. The heuristic strategy is not influenced by the exact payoff values or the agent’s endpoint variance.

These models represent three plausible hypotheses about how people make rapid sensorimotor decisions under risk: (1) people are optimal and maximize expected objective value, (2) people are suboptimal and maximize a biased subjective utility, or (3) people are suboptimal and follow a simple heuristic that works well enough. To test these hypotheses, we generated predictions from the models and compared these predictions to human performance. We first evaluated the models’ absolute goodness of fit, by graphically comparing observed mean reach endpoints with model-predicted mean endpoints. If subjects’ behavior were perfectly captured by a model, then their mean endpoints would be identical to the model-predicted endpoints. We next evaluated the models’ relative goodness of fit by estimating the pointwise likelihood of the data under different models. To compared models, we performed likelihood ratio tests using the Vuong statistic (z_v_) which is the summed log-likelihood ratios (LLRs) between two models, scaled by variance of the LLRs and the sample size (Vuong, 1989). Since this statistic is normally distributed, we infer that one model fits better than another when z_v_ > 1.96 or z_v_ < −1.96, which corresponds to a 95% confidence interval for the null hypothesis z_v_ = 0.

#### Reaching behavior deviated from optimal when financial stakes increased

In terms of absolute goodness of fit, the observed mean endpoints systematically diverged from the model-predicted mean endpoints (Figure 3a,b). In general, people tend to aim closer to the loss region than predicted by the models. This was especially prevalent for the optimal and loss-averse models. The data appeared to be captured better by a perceptual heuristic strategy compared to a utility maximizing strategy.

In terms of relative goodness of fit, the full dataset was more likely under the heuristic model than the optimal model (z_v_ = −7.40) and the loss averse model (z_v_ = 22.7), though the data were more likely under the optimal than the loss averse model (z_v_ = 22.9). The heuristic model had a large advantage over the optimal model when the ratio of loss to gain magnitude was large (z_v_ = −15.4), but when the ratio was small, the optimal model provided a better fit than the heuristic model (z_v_ = 3.82) (Figure 3c). When loss circle was near the gain circle the heuristic model had a large advantage over the optimal model (z_v_ = −7.49), though this advantage was smaller when the circles were more distant (z_v_ = −2.74) (Figure 3d).

We also examined subject-level fits to check whether some subset of the subjects did follow the optimal strategy. No subject’s overall dataset was fit better by the optimal or loss-averse models compared to the heuristic model, but six subjects were fit best by the heuristic model (−8.10 < z_v_ < −1.96), and for the remaining eleven subjects the differences of fit between the heuristic and optimal models were not statistically significant (−1.5 < z_v_ < 0.30). Since the loss-averse model fits always provided the worst relative fit, we focus exclusively on the heuristic and optimal models in what follows.

When the loss ratio was −1:1, nine subjects were fit better by the optimal model than the heuristic model (2.41 < z_v_ < 5.27), one subject was fit better by the heuristic model (z_v_ = −7.80), and the differences of fit were not statistically significant for the remaining four subjects (−1.78 < z_v_ < 1.50) (Figure 3e). When the loss ratio was −5:1, all subjects were fit better by the heuristic model compared to the optimal model (−7.52 < z_v_ < −2.13) (Figure 3e). When the loss region was nearest to the gain region, eight subjects were fit better by the heuristic model compared to the optimal model (−4.19 < z_v_ < −2.12), and the differences of fit were not statistically significant for the remaining nine subjects (−1.55 < z_v_ < 1.51) (Figure 3f). When the loss region was furthest from the gain region, three subjects were fit better by the heuristic model (−7.98 < z_v_ < −2.35) and the differences of fit were not statistically significant for the remaining 14 subjects (−1.02 < z_v_ < 1.89) (Figure 3f). Overall, these results show that group-level predictive advantage of the heuristic model over the optimal model also obtained at the subject level for many subjects. However, the heuristic model provided better fit only when financial stakes were high, consistent with the hypothesis that highly motivated performance depends more on top-down cognitive processing, which is susceptible to cognitive biases.

### Experiment 2

We obtained results in experiment 1 that were inconsistent with behavior reported in prior work. For this reason, we sought to replicate our results in a second experiment. Importantly, the second experiment was a much closer direct replication of the seminal task used by Trommershauser et al. (Spatial Vision, 2003). 16 healthy human subjects performed this incentivized reaching task. Subjects first trained to quickly reach to visuospatial targets for 10 blocks of 30 trials. This training session included loss and gain regions, but the loss value was always zero. During training, time limits were unlimited in block one, 850ms in blocks two through four, and 700 ms in blocks five through ten. We obtained an estimate of subjects reaching variance using their endpoints from blocks five through ten. According to the optimal model, subjects use this estimate of their own variance to decide where to reach, since their variability determines the probabilities of various outcomes conditioned on a candidate aim-point. Subjects returned two days later to perform another 12 blocks of 30 trials, this time with real payoffs. Loss values (0, −100, −500 points) varied block-wise and the gain values were always +100 points. Subjects accumulated points throughout the training and test session and their total points were converted to cash bonuses at a rate of $0.25 per 1000 points. Finally, subjects performed an independent choice task to assess their trait-level loss aversion. The primary dependent measure for our analyses was the horizontal position of subjects’ reaching endpoints.

#### Reaching behavior was related to loss value, loss aversion, and an estimate of reach variance from training

We fit a multilevel gaussian family regression model to subjects’ reach endpoints to test whether people reached further from the loss region when (1) the loss was large, (2) they have high loss-aversion, or (3) they experienced high endpoint variance in training.

We found strong evidence for no effect of loss ratio on reach endpoint, suggesting that our subjects did not reach further from the loss region when the ratio of loss to gain was large (β = −0.00, HDI = [-0.01, 0.00], pd = 1.0). We also found evidence for no effect of training variance on reach endpoint, suggesting that subjects who experienced large endpoint variance in training did not reach further from the loss region compared to subjects who experienced small variance (β = −0.02, HDI = [-0.08, 0.04], pd = 0.67). Lastly, we found weak evidence of an effect of loss aversion on reach endpoint, suggesting oddly enough that subjects with higher loss aversion may have reached closer to the loss region compared to subjects with loser loss aversion (β = - 0.06, HDI = [-0.13, −0.00], pd = 0.94). In sum, participants behavior did not appear to conform to the optimal strategy throughout the task.

#### Reaching behavior was suboptimal on high stakes trials

In terms of absolute goodness of fit, the observed mean endpoints systematically diverged from the model-predicted mean endpoints (Figure 4a,b). In general, people tend to aim closer to the loss region than predicted by the models. This was especially prevalent for the optimal and loss-averse models. The data appeared to be captured better by a perceptual heuristic strategy compared to a utility maximizing strategy.

**Figure 4.**
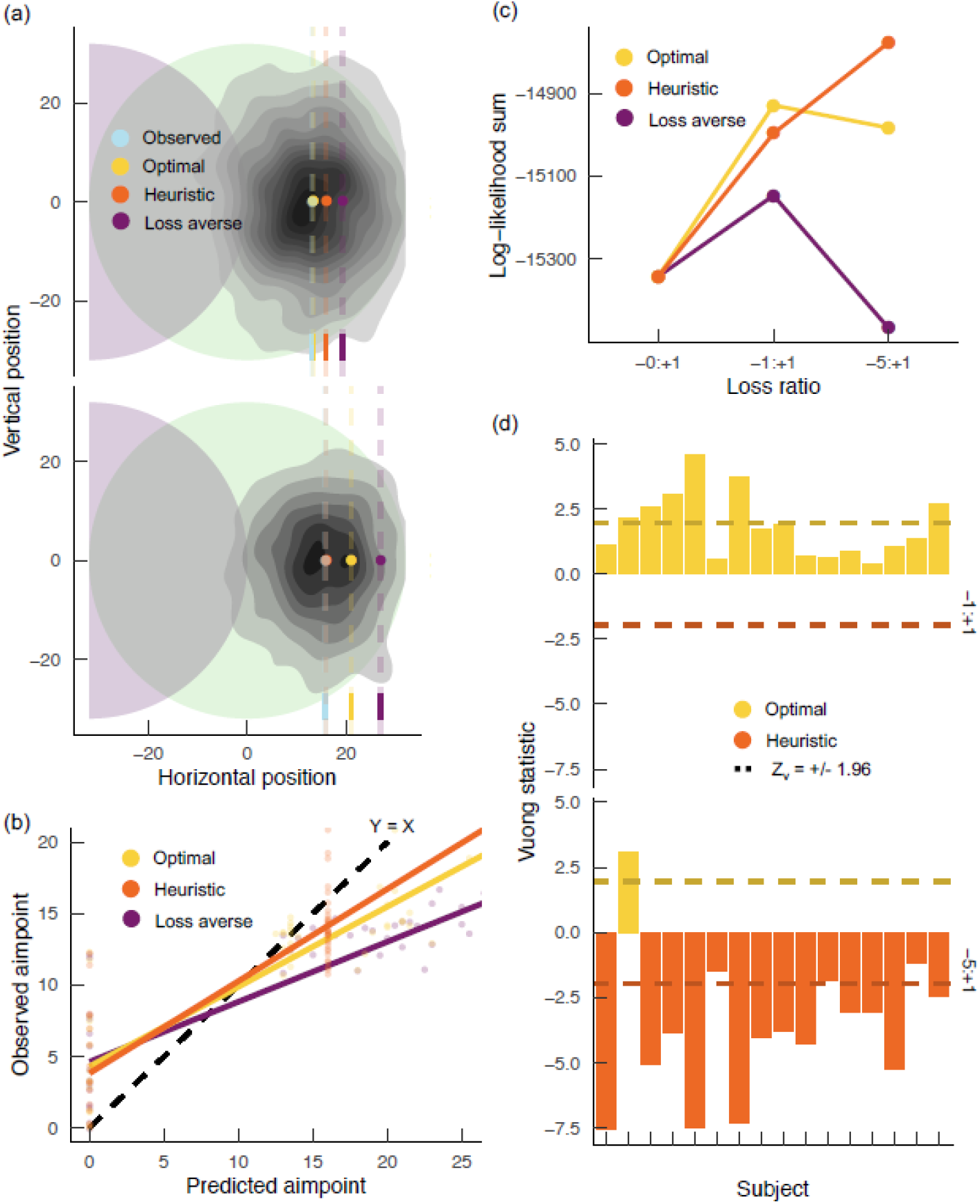
Computational modeling results for Experiment 1. (a) Endpoint distribution and average model predictions for the two loss-ratio conditions. The observed mean endpoint (blue dot) does not move much between the two payoff conditions, similar to the heuristic aimpoint (orange). The optimal aimpoint (yellow) shifts away from the loss region when the loss ratio increases, and this effect is exaggerated for the loss-averse model (purple). This plot is for illustrative purposes, so we show only subjects with above average endpoint variance and above average loss-aversion, to ensure the predictions of the models are visibly distinct. (b) Model-predicted aimpoints vs observed aimpoints. Most people shift less than predicted by the models, but the heuristic predictions provide the best match to the observed data. There is a separate point for each combination of subject, condition and model. (c) Summed log-likelihoods of the data under each model in each payoff condition. The optimal model fits the best in the −1:1 condition by a small amount, and the heuristic model fits best in the −5:1 condition by a greater amount. It also seems that all three models fit worse in in the higher stakes condition. (d) Vuong’s statistics for subject-level heuristic vs optimal comparisons for each payoff condition. The results show that the group-level model-fit patterns hold for most individual subjects.

In terms of relative goodness of fit (Figure 4c), the full dataset was more likely under the heuristic model than the optimal model (z_v_ = −7.37) and the loss averse model (z_v_ = 22.8), though the data were more likely under the optimal than the loss averse model (z_v_ = 25.1). The heuristic model had a large advantage over the optimal model when the ratio of loss to gain magnitude was large (z_v_ = −13.2), but when the ratio was small, the optimal model provided a better fit than the heuristic model (z_v_ = 7.36).

We also examined subject-level fits to check whether some subset of the subjects did follow the optimal strategy (Figure 4d). One subject’s overall dataset was fit best by the optimal model (z_v_ = 3.70), nine subjects were fit best by the heuristic model (−5.57 < z_v_ < −1.96), and for the remaining six subjects, the differences of fit between the heuristic and optimal models were not statistically significant (−1.60 < z_v_ < −0.05). No subject was described better by the loss-averse model than the other models.

When the loss ratio was −1:1, six subjects were fit better by the optimal model than the heuristic model (2.13 < z_v_ < 4.57) and the differences of fit were not statistically significant for the remaining four subjects (0.37< z_v_ < 1.88). When the loss ratio was - 5:1, twelve subjects were fit better by the heuristic model compared to the optimal model (−7.57 < z_v_ < −2.45), one subject was fit better by the optimal model (z_v_ = 3.09), and the differences of fit were not statistically significant for the remaining three subjects (−1.82 < z_v_ < −1.20).

#### Reach variance decreased when financial stakes increased

Lastly, we considered the possibility that our participants respond adaptively to potential losses in ways other than shifting their aimpoint away from the loss region. In particular, we used multilevel Gaussian distributional model to test whether the variance of participants endpoints decreased as the loss ratio increased. Unlike the regression models fit above, this distributional model featured multilevel regression formulae for both the mean and the variance of the outcome variable. The model included fixed effects of loss ratio and random intercepts for each subject.

The model revealed that endpoint variance decreased from the −0:+1 condition to the −1:+1 condition (β = −0.14, HDI = [-0.18, 0.10], pd = 1.0) and from the −1:+1 condition to the −5:+1 condition (β = −0.04, HDI = [-0.08, −0.01], pd = 0.97). This is interesting because endpoint variance limits performance by increasing the probability of outcomes further away from the desired aimpoint, including endpoints in the loss region. Our results suggest that noise in the motor system may be regulated by motivational prospects. However, we examined a variant of the optimal model that used separate variances for each payoff condition, directly estimated from the test data, but this model still provided a worse fit compared to the heuristic model.

### Discussion

We measured sensorimotor decision making using a visually guided reaching task with monetary incentives. This task was of particular interest because it has an “optimal” solution, whereby a person selects reaching aimpoints with maximal expected utility. We tested the hypothesis that people make optimal sensorimotor decisions by comparing human behavior in two experiments to the precise quantitative predictions of optimal and suboptimal computational models. Across both experiments, we found compelling evidence that human behavior deviated systematically from the optimal strategy. The key deviation of human behavior from the ideal was manifest as a tendency to aim too close to the loss region on trials in which financial stakes were high. In these situations, human behavior was captured better by a sub-optimal heuristic strategy that was insensitive to exact payoff values.

According to the optimal model of incentivized reaching behavior, participants use a memory of their endpoint variance in training and an observation of the current potential losses and gains to estimate the expected utility associated with different reach aimpoints. Thus, if participants follow this strategy they will aim further from the loss region when the loss-to-gain ratio is large (−5:1) compared to when it is small (−1:1), and participants with large endpoint variance will aim further from the loss region than participants with small endpoint variance. Across two experiments, Bayesian regression analyses showed that endpoint variance had minimal influence horizontal reach endpoint. This finding was inconsistent with the optimal model of human performance in this task. We therefore formulated a heuristic model of task performance that does not depend directly on endpoint variance. Follow-up model-comparison analyses revealed that reach endpoint data were significantly more likely under the heuristic model compared to the optimal model. Importantly, this advantage was driven by behavior in the large loss ratio condition, where nearly all subjects were fit significantly better by the heuristic model compared to the optimal. When the loss ratio was −1:1, the models had similar fits to the data, but the optimal model had a small advantage overall. These results suggest that participants deviated from the optimal strategy as the loss ratio increased from −1:1 to −5:1.

Prior work has similarly shown that human visuomotor decision making under risk is suboptimal in some variants of the incentivized reaching task. For example, human performance has been shown to be suboptimal with rapidly varying payoff conditions (Neyedli and Welsh, 2014), complex expected value landscapes (Wu et al., 2006), and delayed onset of payoff-information (Trommershauser 2006b). However, much of this prior work assumes that the seminal work using this task provides strong evidence for near-optimal visuomotor decision making under risk. We show that this assumption may be unfounded and that even the original paradigm elicits behavior that deviates from optimal when financial stakes are high. Sensorimotor behavior may be closer to optimal when stakes are lower because the person does not exert much attention or effort to the task, thus allowing the behavior to unfold in an automatic manner. However, when stakes are high, the behavior will be influenced more by top-down cognitive control processes (Botvinick and Braver, 2014; Yee and Braver, 2018). While heightened investments of attention and effort are bound to enhance performance in some ways, this cognitive dependence may also render the behavior vulnerable to cognitive biases and heuristics. While we found no evidence that people followed the loss-averse strategy, we found consistent evidence that people followed the heuristic strategy when the stakes were high.

A key computational difference between the heuristic and optimal strategies is that the heuristic does not depend on computationally costly estimations of expected utility. If the relative costs of relying on the optimal strategy outweigh its relative benefits for performance, then people may not rely on it. Thus, participants in our experiments may have deviated from the optimal strategy when the loss ratio was large because in that condition the costs of the optimal strategy outweighed its benefits. We confirmed using simulations that the extrinsic benefits of the optimal strategy over the heuristic strategy were smaller when the loss ratio was large (≈+0.15 expected gain, assuming a reward of 1) compared to when the loss ratio was small (≈+0.5 expected gain). Therefore, our subjects probably had less incentive to rely on the more costly on-line optimization strategy when the loss ratio was large. This alone may be sufficient to explain why subjects used the inexpensive heuristic when the loss ratio was large. Furthermore, motivational pressure induced by the high financial stakes might consume additional resources, thereby reducing the total amount of resources available for the decision-making task. This would further increase the effective costliness of the optimal strategy, leading a subject to rely on less expensive heuristic strategies.

We find that subjects do not shift their aimpoints in response to payoff information to the extent that is predicted by the optimal model. The optimal strategy incorporates an estimate of the subject’s endpoint variance and assumes that their variance does not change in response to payoff information. But there is evidence that motivation facilitates the reduction of neuronal noise in the motor system (Manohar et al., 2015). Thus, it was plausible that our subjects had smaller endpoint variance when the loss ratio was large, enabling them to aim closer to the loss region without increasing their risk of loss. Follow-up Bayesian analyses revealed that the variance of participants endpoints decreased as the loss ratio increased. The optimal model as formulated previously makes the dubious assumption that endpoint variance is constant, whereas our data show that endpoint variance varies systematically with financial stakes and may indeed be a key method by which people adapt their sensorimotor behavior in response to increasing financial stakes. However, when we examined a version of the optimal model with condition-specific variances estimated from the test session, it still provided a worse overall fit compared to the heuristic model.

There are several questions unanswered by the present study that could be addressed by future work. Data in the low stakes condition were fit best by the optimal model but data in the high stakes condition were fit best by the heuristic model. However, it remains unknown whether this is because people are switching between the optimal and heuristic strategies, or because people are following some other unknown strategy that would generate the pattern of data observed here. Another unanswered question is whether Another issue with this work is that several theoretically distinct models make very similar predictions (typically within a few pixels) and are associated with similar expected gains in the present task. Therefore, it may be inherently difficult to empirically distinguish between plausible alternative models of behavior in this task and future work should explore alternative tasks that are designed specifically to distinguish optimal and plausible alternative suboptimal models.

In sum, we provide evidence across two incentivized reaching experiments (19,080 endpoints) that humans do not make perfectly rational decisions about where to aim their movements when there are large potential losses at stake. An optimal agent would integrate an estimate of its own endpoint variance with payoff and perceptual information to find an aimpoint with maximal utility. The optimal agent therefore aims further from the loss region when its variance estimate increases <u>*or*</u> when the potential loss-to-gain ratio increases. However, our subjects did not display these behavioral patterns to the extent that was predicted by the optimal model. Instead, our data suggest that participants may follow an alternative heuristic strategy when the potential losses are large. According to this model, the agent solves a simplified version of the decision problem, one in which the solution is more or less invariant to endpoint variance and loss-to-gain ratio. Our participants unequivocally deviated from the objectively rational strategy. However, they also did not behave in a manner fit to be described as irrational. Regarding the question of human rationality, our work provides support for a more moderate position according to which people, with their limited computational resources (time, memory, attention, etc.), tend to solve simplified versions of difficult problems, leading to behavior that satisfices, but does not optimize.

## Methods

### Experiment 1

#### Ethics statement

All procedures were approved by the University of Michigan Institutional Review Board of Health Sciences and Behavioral Sciences and the experiment was carried out in accordance with the guidelines of this review board.

#### Participants

23 students (16 female) from the University of Michigan participated in experiment 1. All participants gave written informed consent to participate in the study. All participants were right-handed, free of neurological disorders, and had normal or corrected-to-normal vision. At the end of the experiment, participants were compensated at a rate of $10/hour plus performance bonuses and their outcome from the loss aversion task described below. A participant’s base hourly compensation was not affected by their performance in any task.

#### Apparatus

Participants sat in an immobile chair in a small room in front of a 23” touchscreen computer monitor, and a keyboard positioned such that the spacebar was 21.5 cm away from the screen. We programmed the reaching task using PsychoPy (31) and the loss aversion task using the PyGame (www.pygame.org).

#### Reaching task

Participants performed an incentivized reaching task that has been argued to elicit movement strategies that are near-optimal adaptations to movement variability and payoffs (28). At the start of each trial (Figure 1), the participant was presented with the reward and penalty values for 1,500 ms. Next, the participant looked at a white fixation cross and then pressed and held the space bar with their right index finger when they were ready to begin. Once the space bar was pressed, the fixation cross became blue, and a blue box (88 mm x 88 mm) was presented at the center of the screen for a randomized duration of 400 to 600 ms to indicate the region where the imperative stimulus could appear. Next, overlapping green and red circles (radii = 8.5 mm) appeared at a random location within the blue bounding box and the participant had 1000 ms to reach out and touch the screen.

If a participant touched within the boundary of the green “reward” circle before the time limit (1,000 ms), they earned a reward. If they touched within the boundary of the penalty circle before the time limit, they incurred a penalty. If they touched within the boundaries of both circles before the time limit, they incurred the sum of the penalty and reward. A touch outside both the penalty and reward circles resulted in neither a penalty nor a reward. If a participant completed their reach after the trial time limit had elapsed, they incurred a large penalty (−$5) for that trial and were shown feedback stating, “Too slow”. If a participant released the space bar before the reward and penalty circles appeared, the trial was aborted, and they were shown feedback stating, “Too soon”.

The experiment consisted of training and test sessions completed on two separate days and lasting approximately one hour each. The training session was divided into three parts. In Part 1, which consisted of 12 blocks of 36 trials, participants reached at a single unchanging yellow dot. This provided a distribution of endpoints from which to estimate each participant’s reaching end-point variability. Part 2 consisted of 6 blocks of 36 trials. During Part 2, participants were exposed to six conditions: 3 payoff conditions ([-$0, +$1]; [-$1, +$1]; [-$5, +$1]) with one of two separation distances between the penalty and reward regions (1R; 1.4R, where R = 8.55 mm). Participants were informed that performance in training did not affect their total earnings in the task and was just for practice. Conditions varied randomly on a trial-by-trial basis. Part 3 of training consisted of 2 additional blocks of 36 trials each in which two additional payoff conditions were added: [-$3, +$3] and [-$15, +$5]. These ‘high-stakes trials’ were essential to the comparison between the loss-averse model and the optimal model as the former would predict more drastic shifts away from the penalty region on these trials.

On Day 2, before the test session, participants took a “pre-test” to ensure they understood the task instructions. This pre-test presented a series of hypothetical reaching endpoints for every possible payoff condition and asked the participant how much would be earned for each item. In order to advance to the test phase of the experiment, participants needed to complete the pre-test with 100% accuracy. If participants did not understand an item, we explained it to them. Only one participant failed to complete the test with 100% accuracy on their first attempt. After the pre-test, participants performed 10 blocks of 60 trials each, with all 10 conditions varying randomly on a trial-by-trial basis. At the end of the test session, one trial was selected at random from each block and participants gained and lost bonus earnings based on their performance on these randomly selected trials. After the reaching task was complete, participants performed the loss-aversion task described below.

#### Loss-aversion task

After participants completed the last block of the test session, they completed a gambling task which measured their loss-aversion, understood as the extent to which they weighted losses more than objectively equivalent gains (3,4). In this task, participants were presented with a randomized series of 50/50 gambles. Potential gains ranged from +$10 to +$30, while potential losses ranged from -$5 to -$15. Participants were instructed to evaluate each gamble independently and to accept or reject each of these gambles. They were further instructed to avoid the use of simple decision-making rules, such as a rule to accept all gambles with potential rewards greater than $10. Participants were told that one of the gambles would be randomly selected after completing the task and that if they had accepted the chosen gamble, the participant would win or lose the amount displayed based on the outcome of a computerized coin flip. Participants were instructed that the outcome of the gamble could add to or take away from the bonus they earned from the reaching task.

To calculate a measure of loss aversion from this task, logistic regression was performed with the size of the potential gain and the size of the loss as independent variables and acceptance/rejection as the dependent variable. This analysis was performed separately for each participant. Behavioral loss aversion (λ) was computed as λ = β_loss_/β_gain_ where both β_loss_ and β_gain_ are the unstandardized regression coefficients for the gain and loss variables separately. This analysis makes the simplifying assumption that subjective value is linearly related to objective value for both losses and gains. Eight participants (four in each experiment) had performance that resulted in an implausible λ value that was less than one (loss-seeking) or greater than 15 (extremely loss-averse). These participants appeared to be following a simple decision-making rule rather than evaluating each gamble independently. If λ < 1, we let λ = 1, and if λ > 5, we let λ = 5.

### Experiment 2

#### Ethics Statement

All procedures were approved by the University of Michigan Institutional Review Board of Health Sciences and Behavioral Sciences and the experiment was carried out in accordance with the guidelines of this review board.

#### Participants

18 students (12 female) from the University of Michigan. All participants gave written informed consent to participate in the study. All participants were right-handed and had normal or corrected-to-normal vision. At the end of the experiment, participants were compensated at a rate of $10/hour plus performance bonuses and their outcome from the loss aversion task described below, if applicable. A participant’s base hourly compensation was not affected by their performance in any task.

#### Apparatus

Participants sat in an immobile chair in a small room in front of a 23” touchscreen computer monitor, and a keyboard positioned such that the spacebar was 21.5 cm away from the screen. We programmed the reaching task using OpenSesame (Mathôt et al., 2012) and the loss aversion task using the PyGame (www.pygame.org)

#### Reaching task

Participants performed a task similar to the incentivized reaching task used in Experiment 1. All task parameters were identical except those described below. Experiment 2 consisted of training and test sessions completed on two separate days and lasting approximately one hour each. The training session was composed of 10 blocks of 30 trials each. In training, participants always saw the overlapping reward and penalty regions, but the penalty value was zero. The time-limit for each reaching movement decreased from unlimited in block 1 to 850 ms in blocks 2 – 4 to 700 ms in blocks 5 – 10. The only penalties in the training session were due to reaches completed after the time-limit. The separation distance between the reward and penalty circles was fixed at 1R (32 pixels). We estimated participants’ movement variability from their data in blocks 5 through 10 of training.

The test session was composed of 12 blocks of 30 trials each. In the test session, participants accrued points throughout the experiment that were translated into cash earnings at a rate of 25 cents per 1000 points at the conclusion of the experiment. The reward value was fixed at +100 points and the penalty value was fixed within each block at either zero, −100, or −500 points (4 blocks each, in random order). The time-limit was 700 ms and the penalty for completing a movement after the time-limit was −700 points.

### Statistical modeling

#### Pre-processing

Raw endpoint data were mapped to a standard space with the gain circle centered around the origin and the loss circle offset to the right of the target circle. We next exclude endpoints falling outside of +/− 3 sd of the grand mean in the horizontal or vertical direction, so that subsequent mean and variance estimations were not biased by extreme outliers. Our statistical modeling focused solely on the horizontal dimension of the data, because only this dimension should be systematically related to financial stakes. Finally, we scaled and centered these data to have a grand mean of zero and a grand standard deviation of 0.5 (Gelman, 2008). We transform our continuous predictors (e.g., training variance, loss aversion) in the same manner. We performed all data wrangling using the R package {dplyr}.

#### Multilevel linear mean endpoint model

We first fit multilevel gaussian regression models of horizontal endpoint mean. The model for Experiment 1 included fixed effects of loss ratio, loss multiplier, separation distance, variance from training, and loss aversion, while the model for Experiment 2 only included effects of loss ratio, variance, and loss aversion. Both models included random intercepts for each subject and treated variance as an auxiliary constant parameter. The model can be expressed as:

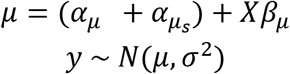

where *y* is an endpoint, *μ* is an endpoint mean, *σ* is a constant group-level endpoint sd, *α* is a group-level intercept term, *α*_*S*_ is a subject-level intercept term, *X* are the predictors (e.g., loss ratio), and *β* are the group-level coefficients for those predictors..

#### Multilevel linear distributional endpoint model

We fit a second multilevel gaussian regression model to the endpoint data from Experiment 2. This model included regression formulae for both the mean and variance of the endpoints. The formulae were the same as those used in the previous models. The model can be expressed as:

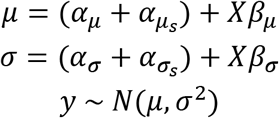

where *y* is an endpoint, *μ* is an endpoint mean determined by the linear equation, *σ* is an endpoint sd determined by the linear equation, *α* is a group-level intercept term, *α*_*s*_ is a subject-level intercept, *X* are the predictors (e.g., loss ratio), and *β* are the group-level coefficients for those predictors.

#### Parameter estimation

We used weakly informative *N*(0,1) priors for all free parameters. This assigns high probability to small estimates and low probability to large estimates because our outcome and predictors have means of zero and standard deviations of 0.5. We estimate the posterior probability distribution over our multidimensional parameter space using No-U-Turns Markov Chain Monte Carlo sampling with 4000 iterations and 50% burn-in. We examine marginal posterior distributions to draw inferences about the size and direction of specific parameter estimates.

Our models were specified and fit using the R package {brms} (Bürkner, 2018, 2017), which compiles probabilistic models in Stan (Carpenter et al., 2017). To quantify the strength of evidence for the median effect size, we use the width of the 89% highest density interval (HDI) (J. Kruschke, 2014). To quantify the strength of evidence for the median effect direction, we use the probability of direction (pd). We estimated these measures using the tidy_stan() function from the R package {sjstats} (Lüdecke, 2019). There were no divergent transitions in sampling, the MCMC traces had normal appearances, the r-hat statistics for all parameters were 1.0, the effective sample sizes were large, and the Markov chain standard errors were approximately zero.

To check the predictive validity of the models, we performed posterior predictive checks using 4000 draws from the estimated posterior. These posterior replications were used to predict observed endpoint densities, means, and variances, and graphical tests revealed a good fit between the predicted and observed summary statistics. We extracted posterior predictions using the function posterior_predict() from the R package {rstantools}. To visualize the predicted effects, we estimated marginal effects of predictors on the outcome using brms::marginal_effects(). All plots were created using the R package {ggplot2} (Wickham, 2016).

### Computational modeling

Linear regression models are useful for detecting simple linear relationships between behavior and predictors of interest, but they give little insight about the precise computations that are performed by the subject to generate their behavior. To address this, we formulated some computational models which embody different hypotheses about the computations used to generate observed reaching behavior in our task.

#### Optimal model

This model is a very slightly modified variant of the MEGaMove model (Trommershäuser et al., 2003). The model states that reaching behavior depends on the estimation and maximization of utility surfaces over the stimulus space. These utility surfaces depend on the perceived geometry of the stimulus, the financial stakes associated with different reach outcomes, and the subject’s internal estimate of their endpoint variance (i.e., outcome uncertainty). The model determines an expected utility for all aimpoints under consideration and the maximum of this surface (i.e., the optimal aimpoint) is given by:

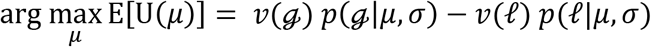

where *μ* is a 2d aimpoint, ℊ is the gain circle, ℓ is the loss circle, *ν*(*c*) returns the value of an endpoint landing in a circle, and *σ* is the variance of endpoints from training. The probability a reach endpoint landing inside an offset circle is given by:

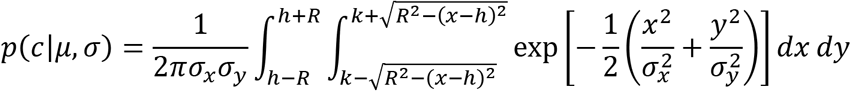

where the aimpoint *μ* = (0,0), *c* is a circle centered around (*h*, *k*) with radius *R* and where the correlation between *x* and *y* is zero (DiDonato and Jarnagin, 1961). In the present study, we did not assume *ρ* = 0, but instead used full variance-covariance matrices estimated from training data, however the simpler case is sufficient to illustrate how the model works. We used the function pmvnEll() from the R package {shotGroups} to numerically solve the above integrals.

#### Loss-averse model

This model is a simple extension of the optimal model. It differs only in that it incorporates an estimate of subjects’ trait-level aversion to financial loss. Loss aversion was estimated in a separate financial choice task in which participants chose whether or not to accept various 50/50 gambles (Tom et al., 2007). Choice data are used to estimate separate linear utility functions over gains and losses. Loss-aversion was defined as the ratio of the slopes of the utility functions for losses compared to gains. In the loss-averse model, losses are multiplied by participants loss aversion when the utility surfaces are estimated. This decreases the utility of aimpoints in proportion to their distance from the loss region and leads to utility maxima that are distinct from the maxima of the optimal utility surfaces, with the loss-averse maxima being further from the loss region compared to the optimal maxima. The loss-averse aimpoint is given by:

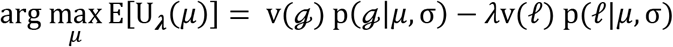

where *μ* is a 2d aimpoint, ℊ is the gain circle, ℓ is the loss circle, v(*c*) returns the value of an endpoint landing in a circle, *σ* is the variance of endpoints from training, and *λ* is a loss-aversion measure.

#### Heuristic model

We hypothesized that people might follow a simple heuristic and only keep track of the stimulus configuration and whether there is a potential loss, ignoring their endpoint variance and the exact loss-to-gain ratio. The heuristic involves two simplifications: (1) ignoring the vertical dimension of the stimuli and (2) binarizing the potential payoffs. This reframing isolates a 1d target (gain) interval with zero-return margins. The goal of the heuristic strategy is to maximize the probability of making contact with this 1d target interval:

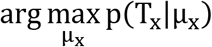

where T is the target interval and the μ_x_ is a candidate aimpoint. Interestingly, the solution to the above goal is invariant to the participant’s endpoint variance and is given by:

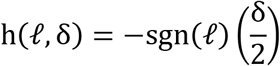

where ℓ is the loss value and δ is the distance between the centers of the loss and gain regions.

The heuristic model is simpler than the previous models in terms of information-processing complexity. For instance, the optimal model gives different predictions for each loss ratio and subject (given unique endpoint variances), while the heuristic model does not. These invariances also make the strategy more robust, since the heuristic strategy is locally optimal for any given endpoint variance (outcome uncertainty), changes in endpoint variance can be ignored. Nonetheless, performance (i.e., expected gain) using the heuristic strategy is enhanced when endpoint variance gets smaller, therefore motivational enhancements to heuristic performance may be mediated by changes in endpoint variance rather than changes in endpoint mean.

#### Model comparison

The models provide quantitative predictions of mean endpoints across subjects and task conditions. The models and predictions were a priori in the sense that they were independent from the test data, with the exception of the heuristic model which was formed after looking at data from Experiment 1. For each observed endpoint and model, we estimated its likelihood given the model-predicted aimpoint (*μ* = (*μ*_*X*_, *μ*_*Y*_)) and the subject’s covariance matrix (σ = [*σ*_*X*_, cov(*X*, *Y*), *σ*_*Y*_, cov(*Y*, *X*)]) estimated from training data. These likelihoods are given by:

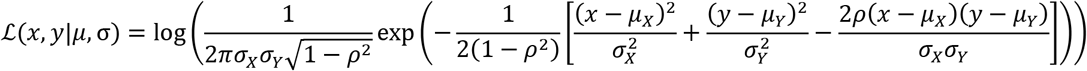

where *ρ* = cov(X, Y)/(*σ*_*X*_*σ*_*Y*_) and represents the correlation between *X* and *Y*. We estimated these likelihoods using the dmvnorm() function from the R package {mvtnorm}.

We then performed likelihood ratio tests between two models to test whether the data were significantly more likely under one model compared to another. Since our models were not nested, we used Vuong’s log-likelihood ratio test to compare their associated likelihoods (Vuong, 1989). And because our models all have no free parameters (*k* = 0), Vuong’s statistic between two models simplifies to:

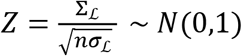

where *n* is the number of observations and:

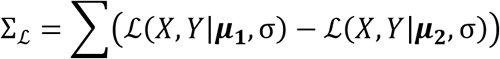

and:

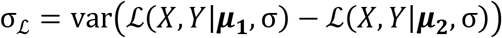

where *X* and *Y* are vectors of observed horizontal and vertical endpoints, *μ*_*N*_ is a vector of predicted aimpoints from a model, and *σ* is a vector of subjects’ variance-covariance matrices estimated from training data. Vuong demonstrated that *Z* is normally distributed, therefore we use a threshold of 1.96 to decide whether to reject the null hypothesis that the likelihood of the data does not vary between models.

## References

Baker CL, Saxe R, Tenenbaum JB. 2011. Bayesian theory of mind: Modeling joint belief-desire attribution. Proceedings of the annual meeting of the cognitive science society 33.

Bastos AM, Usrey MW, Adams RA, Mangun GR, Fries P, Friston KJ. 2012. Canonical Microcircuits for Predictive Coding. Neuron 76:695–711. doi:10.1016/j.neuron.2012.10.038

Botvinick M, Braver T. 2014. Motivation and cognitive control: from behavior to neural mechanism. Annual review of psychology 66:83–113. doi:10.1146/annurev-psych-010814-015044

Braun DA, Nagengast AJ, Wolpert DM. 2011. Risk-Sensitivity in Sensorimotor Control. Frontiers in Human Neuroscience 5:1. doi:10.3389/fnhum.2011.00001

Brown H, Friston K, Bestmann S. 2011. Active Inference, Attention, and Motor Preparation. Frontiers in Psychology 2:218. doi:10.3389/fpsyg.2011.00218

Bürkner P-C. 2018. Advanced Bayesian Multilevel Modeling with the R Package brms. The R Journal 10:395. doi:10.32614/RJ-2018-017

Bürkner P-C. 2017. brms : An R Package for Bayesian Multilevel Models Using Stan. Journal of Statistical Software 80. doi:10.18637/jss.v080.i01

Carpenter B, Gelman A, Hoffman MD, Lee D, Goodrich B, Betancourt M, Brubaker M, Guo J, Li P, Riddell A. 2017. Stan : A Probabilistic Programming Language. Journal of Statistical Software 76. doi:10.18637/jss.v076.i01

Chib VS, Shimojo S, O’Doherty JP. 2014. The Effects of Incentive Framing on Performance Decrements for Large Monetary Outcomes: Behavioral and Neural Mechanisms. The Journal of Neuroscience 34:14833–14844. doi:10.1523/JNEUROSCI.1491-14.2014

Dayan P, Daw ND. 2008. Decision theory, reinforcement learning, and the brain. Cognitive, Affective, & Behavioral Neuroscience 8:429–453. doi:10.3758/CABN.8.4.429

DeCaro MS, Thomas RD, Albert NB, Beilock SL. 2011. Choking under pressure: Multiple routes to skill failure. Journal of Experimental Psychology: General 140:390. doi:10.1037/a0023466

DiDonato A, Jarnagin M. 1961. Integration of the General Bivariate Gaussian Distribution over an Offset Circle. Mathematics of Computation 15:375. doi:10.2307/2003026

Diedrichsen J, Shadmehr R, Ivry RB. 2010. The coordination of movement: optimal feedback control and beyond. Trends in Cognitive Sciences 14:31–39. doi:10.1016/j.tics.2009.11.004

Englert C, Oudejans RR. 2014. Is Choking under Pressure a Consequence of Skill-Focus or Increased Distractibility? Results from a Tennis Serve Task. Psychology 2014:1035–1043. doi:10.4236/psych.2014.59116

Feldman H, Friston KJ. 2010. Attention, Uncertainty, and Free-Energy. Frontiers in Human Neuroscience 4:215. doi:10.3389/fnhum.2010.00215

Friston K. 2011. What Is Optimal about Motor Control? Neuron 72:488–498. doi:10.1016/j.neuron.2011.10.018

Friston K. 2010. The free-energy principle: a unified brain theory? Nature Reviews Neuroscience 11:127. doi:10.1038/nrn2787

Gelman A. 2008. Scaling regression inputs by dividing by two standard deviations. Statistics in Medicine 27:2865–2873. doi:10.1002/sim.3107

Gelman A, Jakulin A, Pittau M, Su Y-S. 2008. A weakly informative default prior distribution for logistic and other regression models. The Annals of Applied Statistics 2:1360–1383. doi:10.1214/08-AOAS191

Gray R. 2004. Attending to the Execution of a Complex Sensorimotor Skill: Expertise Differences, Choking, and Slumps. Journal of Experimental Psychology: Applied 10:42. doi:10.1037/1076-898X.10.1.42

Griffiths TL, Kemp CT, Tenenbaum JB. 2008. Bayesian Models of Cognition.

Howes A, Warren PA, Farmer G, El-Deredy W, Lewis RL. 2016. Why contextual preference reversals maximize expected value. Psychological Review 123:368. doi:10.1037/a0039996

Kinrade NP, Jackson RC, Ashford KJ. 2010. Dispositional reinvestment and skill failure in cognitive and motor tasks. Psychology of Sport and Exercise 11:312–319. doi:10.1016/j.psychsport.2010.02.005

Knill DC, Richards W. 1996. Perception as Bayesian inference, Cambridge University Press. Cambridge University Press.

Körding KP, Wolpert DM. 2006. Bayesian decision theory in sensorimotor control. Trends in Cognitive Sciences 10:319–326. doi:10.1016/j.tics.2006.05.003

Körding KP, Wolpert DM. 2004. Bayesian integration in sensorimotor learning. Nature 427:244. doi:10.1038/nature02169

Kruschke J. 2014. Doing Bayesian Data Analysis: A Tutorial with R, JAGS, and Stan, 2nd ed, Academic press. Academic press.

Kruschke John. 2014. Doing Bayesian data analysis: A tutorial with R, JAGS, and Stan, Academic Press. Academic Press.

Lee TG, Grafton ST. 2015. Out of control: diminished prefrontal activity coincides with impaired motor performance due to choking under pressure. NeuroImage 105:145–55. doi:10.1016/j.neuroimage.2014.10.058

Lewis RL, Howes A, Singh S. 2014. Computational rationality: Linking mechanism and behavior through bounded utility maximization. Topics in cognitive science 6:279–311.

Ma WJ, Beck JM, Latham PE, Pouget A. 2006. Bayesian inference with probabilistic population codes. Nature Neuroscience 9:1432–1438.

Makowski D, Ben-Shachar MS, Chen AS, Lüdecke D. 2019. Indices of Effect Existence and Significance in the Bayesian Framework. Frontiers in psychology 10:2767. doi:10.3389/fpsyg.2019.02767

Manohar SG, Chong T, Apps MA, Batla A, Stamelou M, Jarman PR, Bhatia KP, Husain M. 2015. Reward Pays the Cost of Noise Reduction in Motor and Cognitive Control. Current biology : CB 25:1707–16. doi:10.1016/j.cub.2015.05.038

Neyedli HF, Welsh TN. 2014. People are better at maximizing expected gain in a manual aiming task with rapidly changing probabilities than with rapidly changing payoffs. Journal of Neurophysiology 111:1016–1026. doi:10.1152/jn.00163.2013

Neyedli HF, Welsh TN. 2013. Optimal weighting of costs and probabilities in a risky motor decision-making task requires experience. Journal of Experimental Psychology: Human Perception and Performance 39:638. doi:10.1037/a0030518

Oudejans R, Kuijpers W, Kooijman CC, Bakker FC. 2011. Thoughts and attention of athletes under pressure: skill-focus or performance worries? Anxiety, Stress & Coping 24:59–73. doi:10.1080/10615806.2010.481331

Pezzulo G, Rigoli F, Friston KJ. 2018. Hierarchical Active Inference: A Theory of Motivated Control. Trends in Cognitive Sciences 22:294–306. doi:10.1016/j.tics.2018.01.009

Rahnev D, Denison RN. 2018. Suboptimality in Perceptual Decision Making. The Behavioral and brain sciences 1–107. doi:10.1017/S0140525X18000936

Simon HA. 1955. A behavioral model of rational choice. The quarterly journal of economics. doi:10.2307/1884852

Snyder KM, Logan GD. 2013. Monitoring-induced disruption in skilled typewriting. Journal of Experimental Psychology: Human Perception and Performance 39:1409. doi:10.1037/a0031007

Tauber S, Navarro DJ, Perfors A, Steyvers M. 2017. Bayesian models of cognition revisited: Setting optimality aside and letting data drive psychological theory. Psychological review 124:410–441. doi:10.1037/rev0000052

Tom, Fox C, Trepel C, Science PR. 2007. The neural basis of loss aversion in decision-making under risk.

Trommershäuser J, Gepshtein S, Maloney LT, Landy MS, Banks MS. 2005. Optimal Compensation for Changes in Task-Relevant Movement Variability. The Journal of Neuroscience 25:7169–7178. doi:10.1523/JNEUROSCI.1906-05.2005

Trommershäuser J, Landy MS, Maloney LT. 2006. Humans rapidly estimate expected gain in movement planning. Psychological science 17:981–8. doi:10.1111/j.1467-9280.2006.01816.x

Trommershäuser J, Maloney LT, Landy MS. 2008. Decision making, movement planning and statistical decision theory. Trends in Cognitive Sciences 12:291–297. doi:10.1016/j.tics.2008.04.010

Trommershäuser J, Maloney LT, Landy MS. 2003. Statistical decision theory and the selection of rapid, goal-directed movements. Journal of the Optical Society of America A, Optics, image science, and vision 20:1419–33.

Tversky A, Kahneman D. 1985. The framing of decisions and the psychology of choiceEnvironmental Impact Assessment, Technology Assessment, and Risk Analysis Ssment. Springer. pp. 107–129.

Tversky A, Kahneman D. 1979. Prospect theory: An analysis of decision under risk. Econometrica 47:263–291.

Tversky A, Khaneman D. 1974. Judgment under Uncertainty: Heuristics and Biases. Science 185:1124–1131.

Vuong QH. 1989. Likelihood Ratio Tests for Model Selection and Non-Nested Hypotheses. Econometrica 307. doi:10.2307/1912557

Wickham H. 2016. ggplot2: elegant graphics for data analysis.

Wu S-WW, Trommershäuser J, Maloney LT, Landy MS. 2006. Limits to human movement planning in tasks with asymmetric gain landscapes. Journal of vision 6:53–63. doi:10.1167/6.1.5

Yee DM, Braver TS. 2018. Interactions of motivation and cognitive control. Current Opinion in Behavioral Sciences 19:83–90. doi:10.1016/j.cobeha.2017.11.009

Yu R. 2015. Choking under pressure: the neuropsychological mechanisms of incentive-induced performance decrements. Frontiers in Behavioral Neuroscience 9:19. doi:10.3389/fnbeh.2015.00019

